# Avirulent *Pseudomonas aeruginosa* T3SS-negative strains belonging to Clade 5 produce variable quantities of secondary metabolites

**DOI:** 10.1101/2025.03.27.645720

**Authors:** Selene García-Reyes, Christophe Rusniok, Mylène Robert-Genthon, Eric Faudry, Laura Gomez-Valero, Viviane Chenal-Francisque, Laurent Guyon, Yvan Caspar, Gloria Soberón Chávez, Carmen Buchrieser, Ina Attrée

## Abstract

*Pseudomonas* species are ubiquitous in the environment and serve as valuable source of enzymes and secondary metabolites for industrial applications. *P. aeruginosa* secretes metalloproteases such as elastase LasB and produces bioactive small molecules, including pyocyanin, rhamnolipids, and pyoverdine, with potential biotechnological applications. However, the interest in *P. aeruginosa* for industrial use has been limited due to the virulence-associated Type III Secretion System (T3SS), a key factor in host-pathogen interactions. In this study, we genotypically and phenotypically characterized a collection of *P. aeruginosa* strains naturally lacking T3SS-encoding genes. Phylogenetic analysis revealed that these strains belong to two distinct clades. Several strains exhibited low or no cytotoxicity on epithelial cell lines and were avirulent in the *Galleria* infection model. The level of LasB and the three metabolites — pyocyanin, rhamnolipids, and pyoverdine — varied independently of virulence profiles. Notably, we identified avirulent strains capable of producing at least two secondary metabolites, including monorhamnolipids, highlighting their potential for biotechnological applications.

## Introduction

*Pseudomonas aeruginosa* is a Gram-negative bacterium characterized by its versatile metabolic pathways, intricate regulatory networks, and a wide array of secondary metabolites. Several of these secondary metabolites hold significant biotechnological value, making *P. aeruginosa* an attractive organism for biotechnology applications (Grosjean et al., 2021; Silby et al., 2011) second to *Pseudomonas putida,* a well-established cell factory (de Lorenzo et al., 2024; Weimer et al., 2020). However, these secondary metabolites are also helping *P. aeruginosa* to establish pathogenic interactions with fragilized hosts (Klockgether and Tümmler, 2017).

*P. aeruginosa* synthesizes metalloproteases, exemplified by the secreted elastase LasB (ELA) (Gambello & Iglewski, 1991), and small metabolites such as pyocyanin (PYO) (reviewed in Hall et al., 2016), rhamnolipids (RL) (Pearson et al., 1997) or pyoverdine (PVD) (reviewed in Visca et al., 2007). All of these biomolecules possess biotechnological potential for applications in the biomedical, agriculture, cosmetic, environmental, and pharmaceutical industries.

For example, ELA protease can degrade shrimp waste for chitin preparation and remove the hair from bovine skins and buffalo hides, decreasing the use of sulfide treatment that generates chemical environmental pollution (Ghorbel-Bellaaj et al., 2012). The redox active molecule PYO was proposed to be used in the bioremediation process to improve the crude oil degradation (Norman et al., 2004) or in agriculture for its antifungal activities (DeBritto et al., 2020), and due to its cytotoxic properties, it is also a potential anti-tumor molecule (Abdelaziz et al., 2022; Moayedi et al., 2018).

*P. aeruginosa* synthesizes two types of rhamnolipids (RL) in a coordinated manner alongside several virulence factors: mono-RL, which consist of one rhamnose moiety and a fatty acids dimer, and di-RL, which contains two rhamnose molecules and a fatty acid dimer. These two RL types have different physico-chemical characteristics (Wu et al. 2024); for example, mono-RL has a high capacity to incorporate oil into micelle, applicable for microbial-enhanced oil recovery in the petrochemical industry (Rocha et al., 2020). RL are highly biodegradable and have low toxicity (Soberón-Chávez et al., 2021). Finally, PVD is a siderophore that enables *P. aeruginosa* to grow in low iron environments, but it has affinity for other metal ions like Cd(II), Cu(II), and Ni(II) (Dell’Anno et al., 2022). PVD activity improves the phytoremediation velocity in metal-contaminated sites (Ferret et al., 2015), and can be used to make siderophore-based biosensors and nanosensors for the detection of metals, antibiotics or pesticides (Nosrati et al., 2018). The production of PVD is achieved by the cytoplasmic nonribosomal peptide synthetases PvdL, PvdI, PvdJ, and PvdD, which synthesize the peptide backbone of PVD as well as the components that eventually form the fluorescent chromophore (Lamont & Martin, 2003; Lehoux et al, 2000).

The production of secondary metabolites is regulated by quorum sensing (QS), where at the top of the hierarchy is the Las system, which includes the LasR transcriptional factor and its autoinducer, 3-oxo-dodecanoyl-homoserine lactone (C12-HSL), synthesized by LasI. LasR/3O-C12-HSL positively regulates the synthesis of the Rhl and Pqs systems and ELA. Strains lacking *lasR* do not produce ELA (Pearson et al., 1997). The Rhl system consists of the regulator RhlR and its cognate autoinducer N-butanoyl-L-homoserine lactone (C4-HSL), synthesized by RhlI. This system regulates the production of PYO and RL in *P. aeruginosa* (Brint & Ohman, 1995; Soberón-Chávez et al., 2005, 2021). The third system involves the PqsR transcriptional factor and its autoinducer PQS, synthesized by proteins encoded in the *pqsABCDE* (*pqsA-E*) and *phnAB* operons and the monooxygenase encoded by *pqsH* (Wade et al., 2005). PqsE through its moonlighting chaperon activity (Borgert et al., 2022) further expends RhlR regulon to 389 genes (Letizia et al., 2022)

The use of *P. aeruginosa* in industry has been hampered due to its virulence potential, as well as its high antibiotic resistance. *P. aeruginosa* is considered a major opportunistic human pathogen mainly infecting people with a poor immune response, for example, individuals suffering from cancer, HIV, diabetes, intensive care patients, transplanted patients, with burns or people with cystic fibrosis (Murphy et al., 2008; Reynolds & Kollef, 2021; Sonmezer et al., 2016). Due to the ability of *P. aeruginosa* to evade the action of most antibiotics, the Infectious Diseases Society of America classified it within the “ESKAPE” bacteria with hospital importance (Liu et al., 2021; Pendleton et al., 2013), and the World Organization Health (WHO) classified it as a high research priority for public health (WHO, 2024).

Phylogenetic analyses of the species *P. aeruginosa* defined five phylogenetic groups (Freschi et al., 2019, Ozer et al., 2019). The most populated and studied are groups 1 and 2, represented by the reference strains PAO1 and PA14, respectively. Group 1 and group 2 strains have been isolated from a variety of infections and have a wide geographical distribution; they display cytotoxicity and virulence, notably due to four exotoxins they encode, ExoT, ExoY, and ExoS or ExoU, which are mutually exclusive (Ozer et al., 2019; Pirnay et al., 2009) These exotoxins are exported by a complex secretion nanomachinery, named type III secretion system (T3SS), assembled by >20 macromolecules encoded within the 35kb genomic locus (reviewed in Hauser, 2009).

Interestingly, the entire T3SS locus is missing in strains of the two minor clades, 3 and 5, which are represented by strains PA7 and PA39, respectively (Elsen et al., 2014; Quiroz-Morales et al., 2022; Reboud et al., 2016; Roy et al., 2010). Strains of clade 3 have been referred as “outliers” and were recently reclassified as *P. paraeruginosa* (Rudra et al., 2022). The initial cytotoxic and virulence assays suggested that some strains of clades 3 and 5 might be avirulent, despite the presence of the *exlBA* operon, encoding the pore-forming exotoxin ExlA (Medina-Rojas et al., 2020; Reboud et al., 2016; Quiroz-Morales et al., 2023)

In addition to the absence of T3SS-encoding genes, those strains display other genotypic differences that may influence the synthesis of some secondary metabolites. For example, two genomic regions -*rhlC and phzH-* have been reported to be deleted in some PA7-like strains (Quiroz-Morales et al., 2022) *rhlC* encodes the rhamnosyl transferase RhlC that converts mono-RL to di-RL. *phzH* encodes PhzH, an enzyme involved in the phenazine-1-carboxamide synthesis within the PYO biosynthetic pathway (Mavrodi et al., 2001; Medina et al., 2003; Soberón-Chávez et al., 2005)). Also, deletions in genes encoding five transcriptional regulators have been reported (García-Reyes et al., 2020; Roy et al., 2010).

The systematic impact of these genotypes on the production of secondary metabolites has not been yet investigated, although individual strains from clade 3 have been tested for mono-RL and PYO production (Grosso-Becerra et al., 2016; Sood et al., 2020).

In this work, we genotypically and phenotypically explored a collection of *exlA*+, *T3SS*-negative strains to assess their cytotoxicity, virulence potential, and their susceptibility to antibiotics. We found that the production of three major secondary metabolites PYO, RL, PVD, and the secreted elastase, ELA in five strains belonging to phylogenetic group 5 is variable and independent of their virulence profile. We identified several avirulent strains that maintain their capacity to produce at least two secondary metabolites, including mono-RL, thus representing an opportunity to explore them for biotechnology applications.

## Materials and methods

### Strains and culture conditions

The strains used in this work are listed in the Table 1. Strains were first verified by PCR using following primers and conditions.

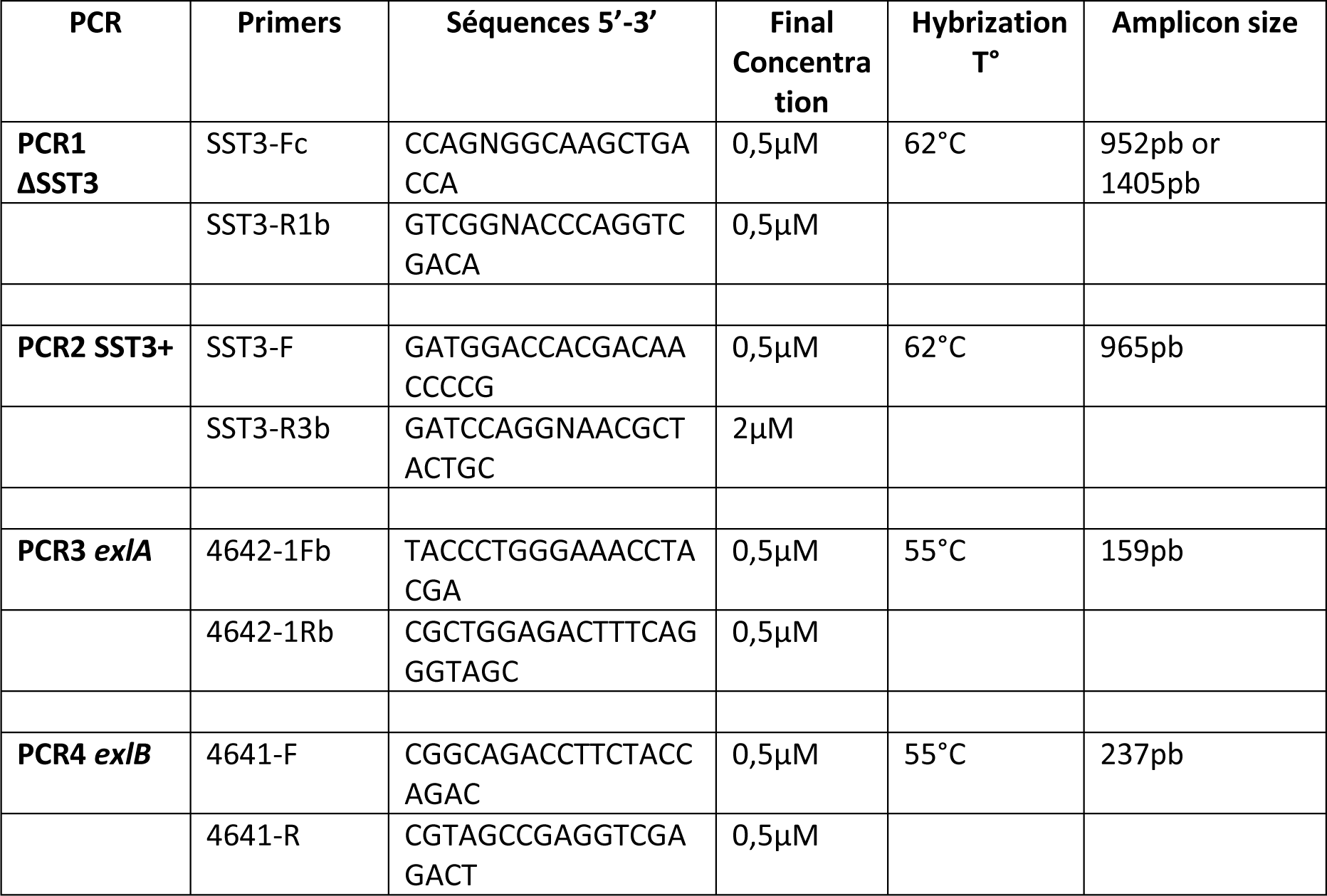

**Table 1.**
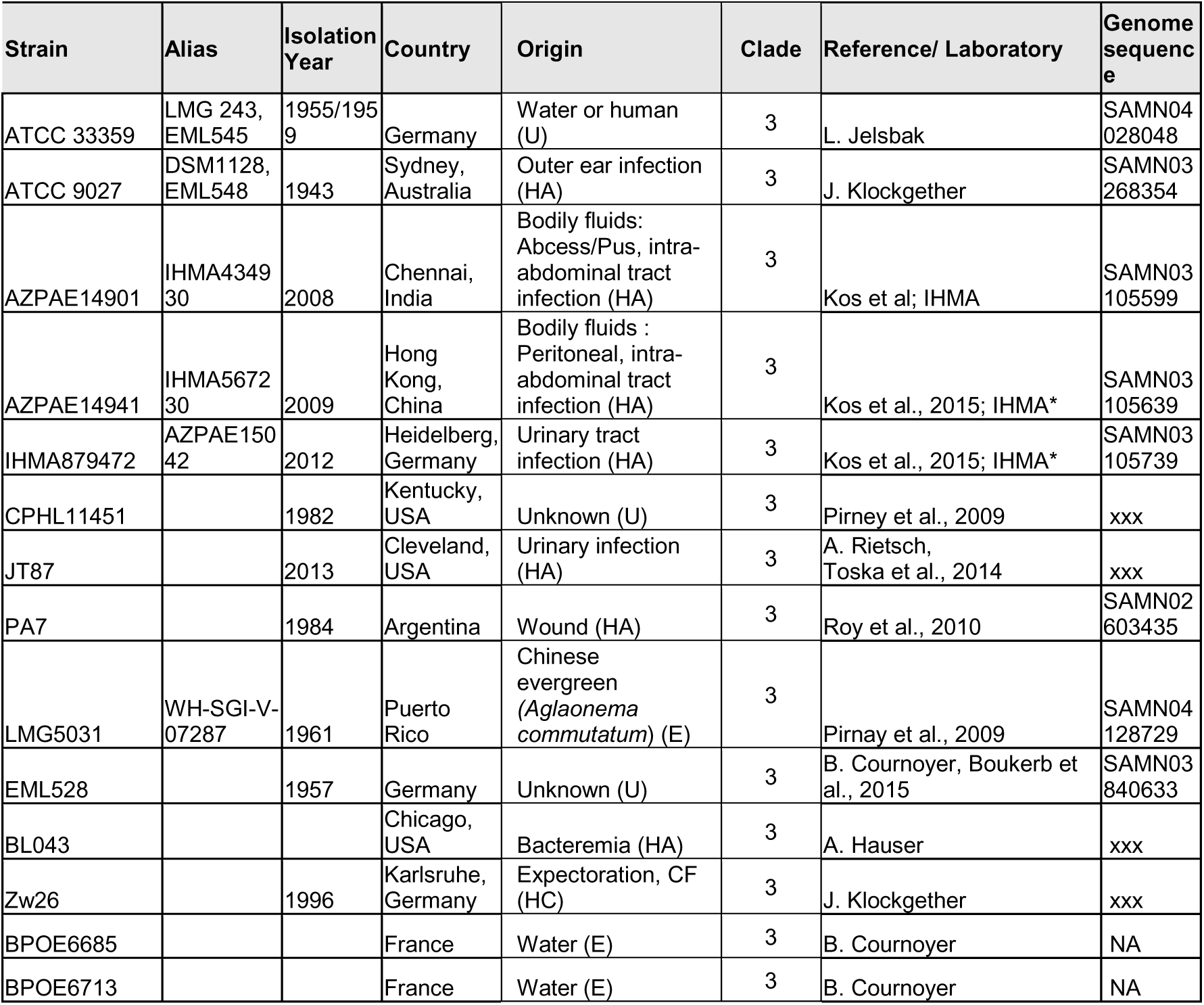
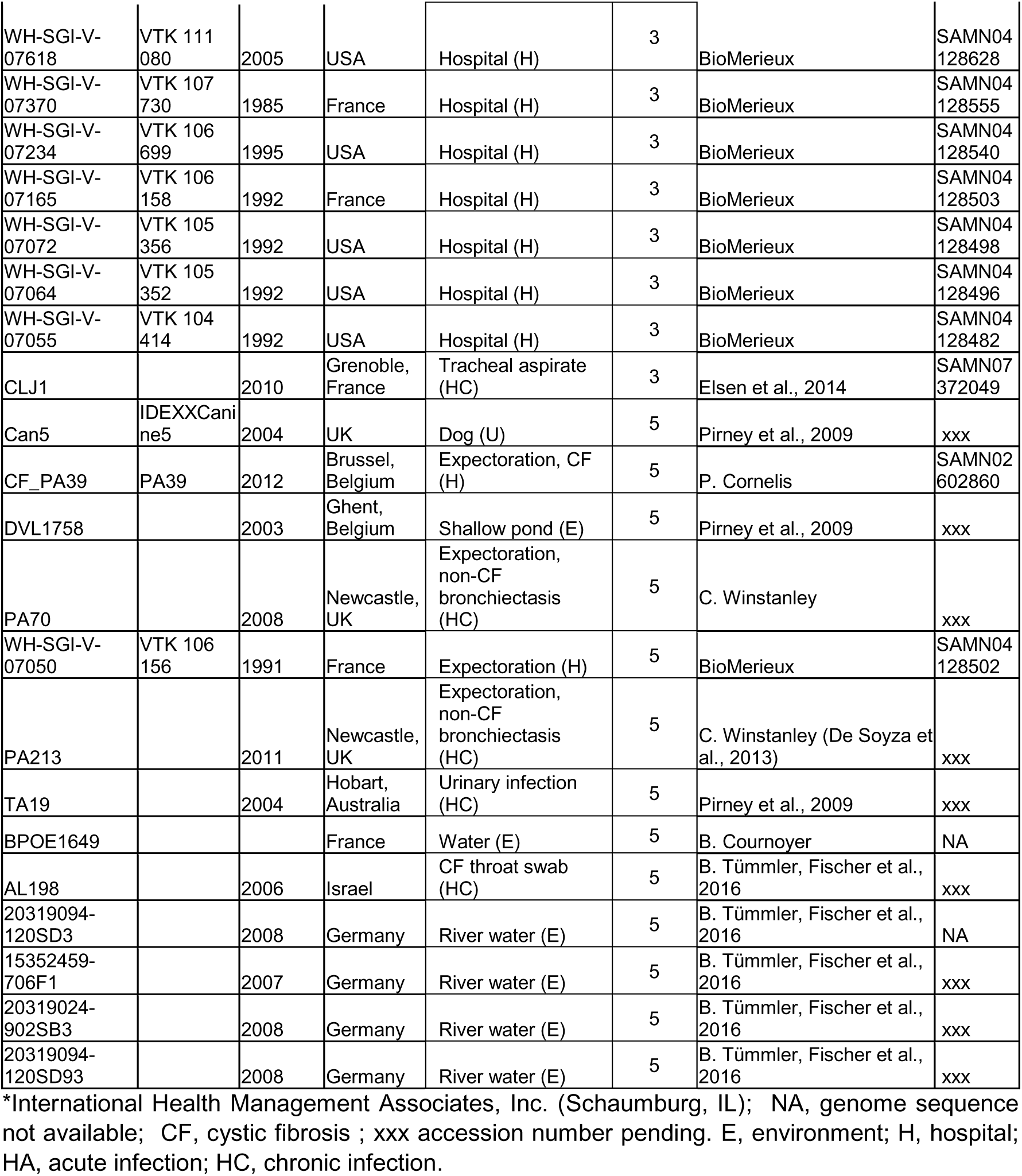
Collection of P. aeruginosa strains belonging to phylogenetic clades 3 and 5 according to Frescki et al., 2015.

| Strain | Alias | Isolation Year | Country | Origin | Clade | Reference/ Laboratory | Genome sequence |
| --- | --- | --- | --- | --- | --- | --- | --- |
| ATCC 33359 | LMG 243, EML545 | 1955/1959 | Germany | Water or human (U) | 3 | L. Jelsbak | SAMN04 028048 |
| ATCC 9027 | DSM1128, EML548 | 1943 | Sydney, Australia | Outer ear infection (HA) | 3 | J. Klockgether | SAMN03 268354 |
| AZPAE14901 | IHMA434930 | 2008 | Chennai, India | Bodily fluids: Abscess/Pus, intra-abdominal tract infection (HA) | 3 | Kos et al; IHMA | SAMN03 105599 |
| AZPAE14941 | IHMA567230 | 2009 | Hong Kong, China | Bodily fluids : Peritoneal, intra-abdominal tract infection (HA) | 3 | Kos et al., 2015; IHMA* | SAMN03 105639 |
| IHMA879472 | AZPAE15042 | 2012 | Heidelberg, Germany | Urinary tract infection (HA) | 3 | Kos et al., 2015; IHMA* | SAMN03 105739 |
| CPHL11451 |  | 1982 | Kentucky, USA | Unknown (U) | 3 | Pirney et al., 2009 | xxx |
| JT87 |  | 2013 | Cleveland, USA | Urinary infection (HA) | 3 | A. Rietsch, Toska et al., 2014 | xxx |
| PA7 |  | 1984 | Argentina | Wound (HA) | 3 | Roy et al., 2010 | SAMN02 603435 |
| LMG5031 | WH-SGI-V-07287 | 1961 | Puerto Rico | Chinese evergreen (*Aglaonema commutatum*) (E) | 3 | Pirnay et al., 2009 | SAMN04 128729 |
| EML528 |  | 1957 | Germany | Unknown (U) | 3 | B. Cournoyer, Boukerb et al., 2015 | SAMN03 840633 |
| BL043 |  |  | Chicago, USA | Bacteremia (HA) | 3 | A. Hauser | xxx |
| Zw26 |  | 1996 | Karlsruhe, Germany | Expectoration, CF (HC) | 3 | J. Klockgether | xxx |
| BPOE6685 |  |  | France | Water (E) | 3 | B. Cournoyer | NA |
| BPOE6713 |  |  | France | Water (E) | 3 | B. Cournoyer | NA |
| WH-SGI-V-07618 | VTk 111 080 | 2005 | USA | Hospital (H) | 3 | BioMerieux | SAMN04 128628 |
| WH-SGI-V-07370 | VTk 107 730 | 1985 | France | Hospital (H) | 3 | BioMerieux | SAMN04 128555 |
| WH-SGI-V-07234 | VTk 106 699 | 1995 | USA | Hospital (H) | 3 | BioMerieux | SAMN04 128540 |
| WH-SGI-V-07165 | VTk 106 158 | 1992 | France | Hospital (H) | 3 | BioMerieux | SAMN04 128503 |
| WH-SGI-V-07072 | VTk 105 356 | 1992 | USA | Hospital (H) | 3 | BioMerieux | SAMN04 128498 |
| WH-SGI-V-07064 | VTk 105 352 | 1992 | USA | Hospital (H) | 3 | BioMerieux | SAMN04 128496 |
| WH-SGI-V-07055 | VTk 104 414 | 1992 | USA | Hospital (H) | 3 | BioMerieux | SAMN04 128482 |
| CLJ1 |  | 2010 | Grenoble, France | Tracheal aspirate (HC) | 3 | Elsen et al., 2014 | SAMN07 372049 |
| Can5 | IDEXXCanine5 | 2004 | UK | Dog (U) | 5 | Pirney et al., 2009 | xxx |
| CF_PA39 | PA39 | 2012 | Brussel, Belgium | Expectoration, CF (H) | 5 | P. Cornelis | SAMN02 602860 |
| DVL1758 |  | 2003 | Ghent, Belgium | Shallow pond (E) | 5 | Pirney et al., 2009 | xxx |
| PA70 |  | 2008 | Newcastle, UK | Expectoration, non-CF bronchiectasis (HC) | 5 | C. Winstanley | xxx |
| WH-SGI-V-07050 | VTk 106 156 | 1991 | France | Expectoration (H) | 5 | BioMerieux | SAMN04 128502 |
| PA213 |  | 2011 | Newcastle, UK | Expectoration, non-CF bronchiectasis (HC) | 5 | C. Winstanley (De Soyza et al., 2013) | xxx |
| TA19 |  | 2004 | Hobart, Australia | Urinary infection (HC) | 5 | Pirney et al., 2009 | xxx |
| BPOE1649 |  |  | France | Water (E) | 5 | B. Cournoyer | NA |
| AL198 |  | 2006 | Israel | CF throat swab (HC) | 5 | B. Tümmler, Fischer et al., 2016 | xxx |
| 20319094-120SD3 |  | 2008 | Germany | River water (E) | 5 | B. Tümmler, Fischer et al., 2016 | NA |
| 15352459-706F1 |  | 2007 | Germany | River water (E) | 5 | B. Tümmler, Fischer et al., 2016 | xxx |
| 20319024-902SB3 |  | 2008 | Germany | River water (E) | 5 | B. Tümmler, Fischer et al., 2016 | xxx |
| 20319094-120SD93 |  | 2008 | Germany | River water (E) | 5 | B. Tümmler, Fischer et al., 2016 | xxx |

\*International Health Management Associates, Inc. (Schaumburg, IL); NA, genome sequence not available; CF, cystic fibrosis ; xxx accession number pending. E, environment; H, hospital; HA, acute infection; HC, chronic infection.

For production of elastase LasB and secondary metabolites, *P. aeruginosa* PAO1 was used as a reference strain. *P. aeruginosa* Δ*lasRrhlR* mutant was used as negative control, when indicated. If not otherwise indicated, the bacterial cultures were prepared in LB media at 37 °C with shaking at 300 rpm.

### Whole genome sequencing

Whole genome sequencing was performed at the Biopolymers Facility, Harvard Medical School, Boston, USA (for strains PA70, DVL1758, JT87, Can5, CPHL11451 and LMG5031) and at Institut Pasteur, Paris, France (strains TA19, PA213, AL198, Zw26, 120SD93, 706F1, 902SB3 and BL043), using the Illumina technology.

### Sequencing and Preprocessing

FastQ files containing sequencing reads were processed in two steps to remove adapter sequences and low-quality regions. First, the Cutadapt software (version 1.15) was used to remove Illumina adapter sequences. Second, Sickle (version 1.33) with a Phred quality score threshold of 20 was employed to trim low-quality bases from the 3’ and 5’ ends of the reads. Additionally, reads shorter than 50 nucleotides after trimming were discarded to ensure sufficient sequence length for genome assembly.

### Genome Assembly

The Spades assembler (version 3.13.1) was used to perform *de novo* genome assembly. Different K-mer values ranging from 21 to 61 were used and combined to optimize the assembly process. Contigs generated by Spades were reordered using CONTIGuator2 software with sequence from different strains of *P. aeruginosa* already sequenced used as references.

### Scaffold Reordering and Annotation

The reordered contigs, forming larger scaffolds, were then annotated using Prokka, a software pipeline embedded within the “annotate” module of PanACoTA (version 1.4.1)

### Pangenome Analysis

The PanACoTA’s “pangenome” module was used to identify the pangenome, which represents the complete set of genes present across all the strains included in the analysis. Protein sequences were clustered into families based on a minimum percent identity threshold of 80% using the “-i 0.8” option.

### Phylogenetic Analysis

The core genome, which consists of genes found in all analyzed strains, was extracted from the pangenome using the PanACoTA’s “corepers” module. Subsequently, the core gene sequences were aligned with the “align” module within PanACoTA. Finally, a phylogenetic tree was constructed using the “tree” module. As parameters, we chose to use the IQtree2 software with the Generalized Time Reversible (GTR) substitution model.

### Analysis of Gene Presence/Absence

To screen the individual genomes for the presence or absence of specific genes or genomic regions, we used orthologous gene families previously established during the core/pangenome analysis. Each gene of interest has been first linked to the corresponding family. Then, we identified, for each strain studied, if the gene of interest has an orthologous. To visualize the presence/absence matrix, we used the Interactive Tree Of Life software (iTOL, https://itol.embl.de)

### Cytotoxicity assay

Epithelial cells (A549, ATCC CCL-185^TM^) were grown in 96-well plates (25×10^3^ cells/well) in 160µl DMEM. In the beginning of the experiment, 40 µl of bacteria at 6×10^8^ bacteria/ml were added corresponding to multiplicity of infection (MOI) of 10, in presence of propidium iodide, 1µg/ml. Propidium iodide incorporation into the cells was followed by the fluorescence emitted at 590nm, every 10 minutes for a period of up to 10 hours, at 37°C, using a Fluoroskan Ascent FL2.5 Microplate Fluorometer (Thermo Corporation) Fluorescence intensities from 100 to 600 min were integrated to calculate Area Under the Curve (AUC) that were divided by the AUC obtained with the IHMA87 strain to give normalized AUCs (AUCnorm) ranging from 0 to 1.

### Antibiotic susceptibility testing

Antibiotic susceptibility testing (AST) was performed using disk diffusion method according to CASFM-EUCAST (Comité de l’antibiogramme de la Société Française de Microbiologie - European Committee on Antimicrobial Susceptibility Testing) guidelines. Briefly a 0.5 Mc Farland bacterial suspension was prepared from bacterial colonies grown overnight at 35°C on tryptic soy agar media. Bacterial suspension was swabbed on a square Mueller Hinton agar plate (MHE agar, BioMérieux, Marcy l’Etoile, France) and antibiotic disks (Ticarcillin 75 μg, Ticarcillin + clavulanic acid 85 μg, Piperacillin 30 μg, Piperacillin + tazobactam 36 μg, Ceftazidime 10 μg, Cefepim 30 μg, Aztreonam 30 μg, Imipenem 10 μg, Meropenem 10 μg, Tobramycin 10 μg, Gentamicin 10 μg, Amikacin 30 μg, Ciprofloxacin 5 μg and Levofloxacin 5 μg ; Bio-Rad, Marnes-la-Coquette, France) were placed on the agar. Plates were incubated under ambient atmosphere at 35 ± 2 °C and zone diameters were measured after 20 ± 4 h of incubation by digital reading (Adagio automaton, Bio-Rad) with manual adjustment of zone diameters. Results were interpreted using CASFM-EUCAST 2023 breakpoints. The strains were classified as sensitive to all tested antibiotics (S), non-multidrug resistant (non-MDR), MDR and extensively (X)-DR using the definitions of European Centre for Disease Prevention and Control (Magiorakos et al., 2012).

### Infections of *Galleria mellonella* larvae

The wax moth *Galleria mellonella* larvae were purchased from Sud-Est Appats (http://www.sudestappats.fr). Uniformly white larvae measuring around 3 cm were further selected upon reception. The bacteria were grown in tubes in LB with shaking to an optical density measured at 600 nm (OD_600_) of 1 and then diluted in PBS to 1000 bacteria/ml. Bacterial solutions were loaded to sterile insulin cartridges. 10 μl of bacterial suspensions were injected into the larvae using an insulin pen. The exact number of bacteria was determined by spotting aliquots with the same pen onto agar plates and counting colonies after growth at 37 °C for 16 h. Injected *Galleria* were maintained in Petri dishes at 37 °C. Dead larvae were counted over 24h. Twenty larvae were used per condition, and the experiment was performed at least twice for each strain.

### Determination of PYO, PVD, ELA and RL production

To analyze PYO production, cultures were diluted to an OD_600_ of 0.01 and incubated in 30 mL LB medium for 16 h with shaking (225 rpm) at 37 °C. PYO was extracted with chloroform and measured by spectrophotometry at 520 nm in acidic solution, as reported previously (Essar et al., 1990). PVD was measure in the culture supernatants by spectrophotometry at 405 nm according to Diggle et al. 2007. The ELA was measure using elastin-Congo Red as substrate (Sigma Aldrich). Briefly, overnight LB broth with antibiotic cultures of each strain were diluted to an OD_600_ of 0.1 and grown for 16 hours in LB media without antibiotic at 37°C with shaking. Then, the cultures were centrifuged (14,000 x g for 5 minutes) and 100 μl of each supernatant was added to Eppendorf tubes containing 5 mg of elastin-Congo Red in 900 μl of 0.1 M Tris-Cl pH 7.2, 1 mM CaCl_2_. Tubes were shaken (225 rpm) at 37 °C for 3 hours. The insoluble substrate (unreacted elastin-Congo red) was pelleted by centrifugation (13,000 x g for 10 minutes) and the absorbance of the supernatant was measured at 495 nm.

The RL were determined by thin layer chromatography (TLC) according to the technique reported (González-Valdez et al., 2024). Briefly, the supernatants were obtained by centrifugation 14,000 x g for 10 minutes at 4°C. 5 mL of each supernatant was be placed in 50 mL Falcon tubes, and then the extraction was carried out adding to each tube 20 µL of HCl and 5 mL of the 2:1 mixture of chloroform and methanol. The organic phase was recovered from each tube and allowed to evaporate at room temperature in an extraction hood. 100 µL of methanol was added to the dry tubes and 5 µL of each sample was loaded onto the TLC plates (TLC SILICA GEL 60 F254, Merck). The TLC was developed using the mobile phase chloroform-methanol-acetic acid, 65:15:2. Finally, the plate was sprayed with a solution of α-naphthol and heated in the oven at 80°C.

### Data analysis

Data from cytotoxicity and larvae infections were processed, plotted and submitted to statistical tests with R version 4.3.2 (R Core Team, 2023). Larvae survival kinetics were analyzed in R using the survival (Therneau, 2024) and gsurvfit packages (Sjoberg et al., 2024).

### Statistical analysis

The statistical analyses were performed using the Graph Pad Prism 9.4.1 software for Analysis of variance (ANOVA) and multiple comparison analysis test (Tukey) to determine the significance of the differences between the means. The values of P ≤ .05 were considered statistically significant.

## Results and discussion

### Establishment of a collection of *T3SS* negative *P. aeruginosa* strains

The strains for this study were collected from different sources (Table 1) based on the absence of the T3SS-encoding locus and the presence of the *exlA* gene (Elsen et al., 2014; Reboud et al., 2016). Upon the reception, the strains were PCR-verified and plated on the *Pseudomonas* Isolation Agar medium. The over-night plate-grown culture was recovered and frozen in duplicates or triplicates on sterile beads. One bead was used for each experiment. Based on the size of the deletion scar obtained by PCR at the T3SS locus (Elsen et al., 2014), the strains were classified either in the clade 3 or 5, according to Freschi et al., 2019. Altogether, the collection contains 35 strains, with 23 and 12 strains belonging to clade 3 and 5, respectively. Nine strains are environmental isolates, and the rest are from human infections, either chronic or acute. Two strains, WH-SGI-V-07050 and Can5, for which we had no information about the exact origin, were classified in “hospital” and “unknown”, respectively. The collection contains strains from Europe (France, Germany, Belgium, UK), South America (Puerto Rico and Argentina), Australia, Israel, India, China and USA.

To understand the genomic diversity of the collection, the genomes of fourteen strains of the collection were sequenced and compared to genomes of *exlBA*^+^ strains available at NCBI. The assemblies were submitted to GenBank (accession numbers pending). For characterization of the strains with respect to the origin, the date of isolation, the source and the genome accession numbers see Table 1.

### Presence/absence of virulence genes correlates with the size of the T3SS scar and the phylogenetic grouping of the strains

Genome analyses confirmed the presence of the *exlBA* operon and the absence of the T3SS-encoding operons as well as the absence of all four effector-encoding genes. The phylogenetic tree of these strains, based on the core genome, groups them in the two previously identified clades (Figure 1). The deletions in the regions around *phzH* and *rhlC* genes, were detected in all strains from our collection. Interestingly, and similarly to the scars of the T3SS region which is variable in size between the two clades (Elsen et al., 2014), the strains could be further differentiated by a deletion of 5 genes (*fosA*, *rhlC*, *PA1131*, *PA1132* and *PA1133*) in clade 3, whereas strains of clade 5 lack only *rhlA* and *PA1133*. Absence or presence of these genes are given in Figure 1 and clearly correlate with the phylogenetic division in clade 3 or clade 5. We also analysed the presence of genes encoding major regulators. Whereas some strains are lacking some regulatory genes (i.e. *lasR*, *pqsR*) as has been previously suggested (Quiroz-Morales et al., 2023), we did not identify any common signature in regulatory genes between the strains of the two clades.

**Figure 1.**
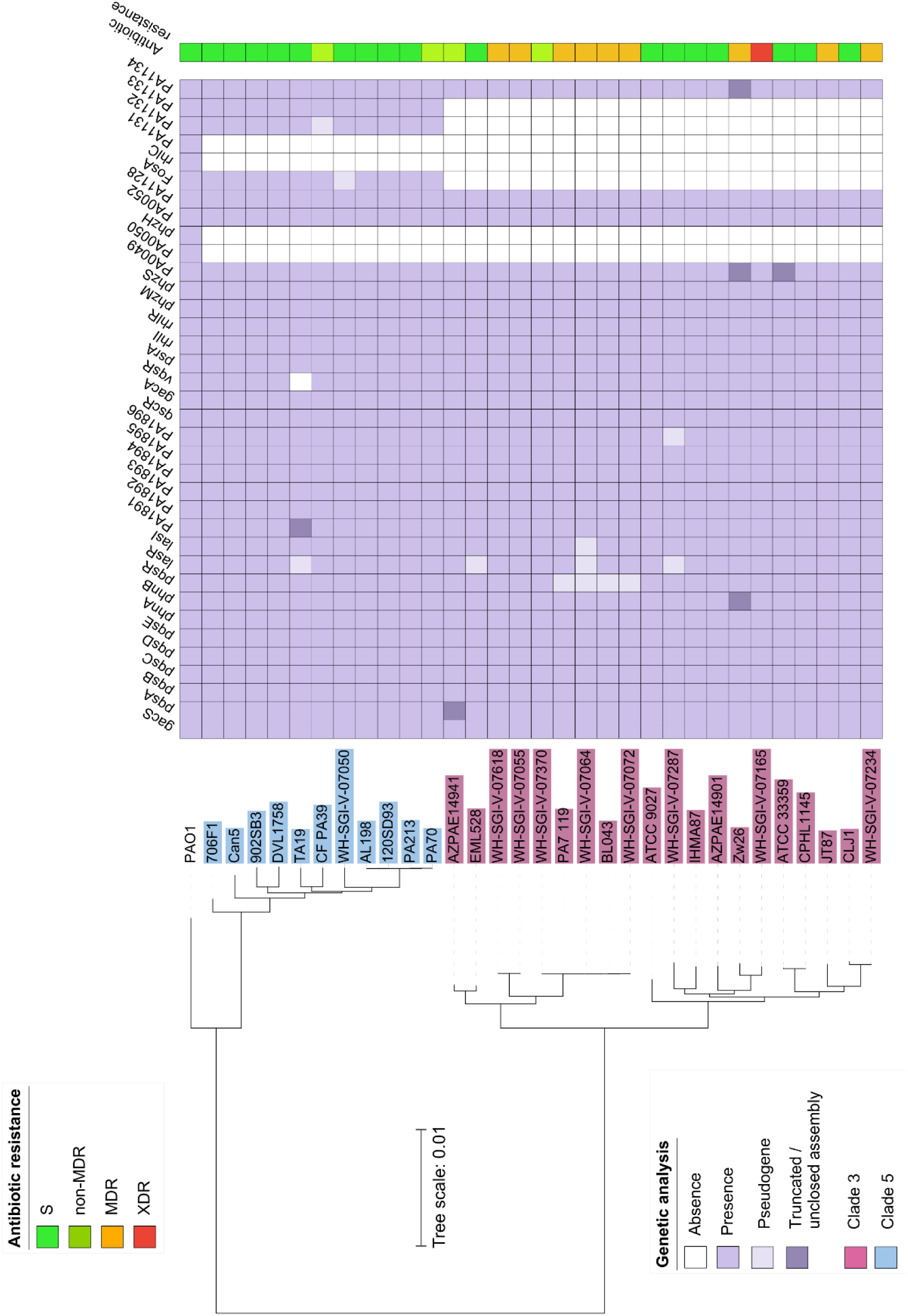
The T3SS-negative, *exlBA^+^* strains cluster in two distinct phylogenetic groups. - clades 3 and 5-. Reference strain PAO1 is representative of Clade 1. Based on the whole-genome analysis, the clade 3 strains were reclassified as a new species *P. paraeruginosa* (Rudra et al., 2022). The genomes were analysed for presence/absence of several genes or genomic regions (shown as cells colored in purple/white). Pseudogene (in light purple): the sequence is present, but contains several mutations, including stop codons. Truncated / unclosed assembly (in darker purple): the complete gene could not be detected due to the unclosed assembly. The resistance of individual strains (Table S1) is reported in a separate line, and indicated as sensible (S, green), non-<u>m</u>ulti<u>d</u>rug resistant (non-MDR, yellow), MDR (orange) and XDR (red).

### Strains belonging to Clade 5 are globally less cytotoxic on epithelial A549 cells

The cytotoxic potential of all the strains present in the collection was assessed on the epithelial A549 cells incubated with bacteria in presence of propidium iodide (PI). The kinetics of the PI incorporation into the cells, due to the pore forming activity of ExlA, was measured over a 10h-period on a microplate fluorometer. The Area Under the Curve (AUC) of each kinetics was calculated as a proxy for cytotoxicity: the faster the cells die, the higher is the AUC as described before (Basso et al., 2017). The AUCs from each independent experiment were normalized against the ones obtained with the fully cytotoxic ExlA-positive strain IHMA87 (Basso et al., 2017; Elsen et al., 2014).

The collection showed high variability in the cytotoxicity profiles, ranging from highly cytotoxic strains reaching 100% cell death within 3h post-infection (IHMA87, AZPAE14941, EML528 or BL043) to strains displaying no cytotoxicity - PA213, TA19, PA70-(Figure S1). Medians of the AUCs of each strain were grouped according to their clades or the origins and the comparison showed significant difference between the two clades, with clade 5 being less cytotoxic (t-test p-value = 0.00027) (Figure 2A). Significant difference was observed when examining the origins of the strains (Figure 2B) with strains isolated from chronic infections exhibiting less cytotoxicity. Globally, strains obtained from the environment or acute hospital infections showed higher cytotoxicity.

**Figure 2.**
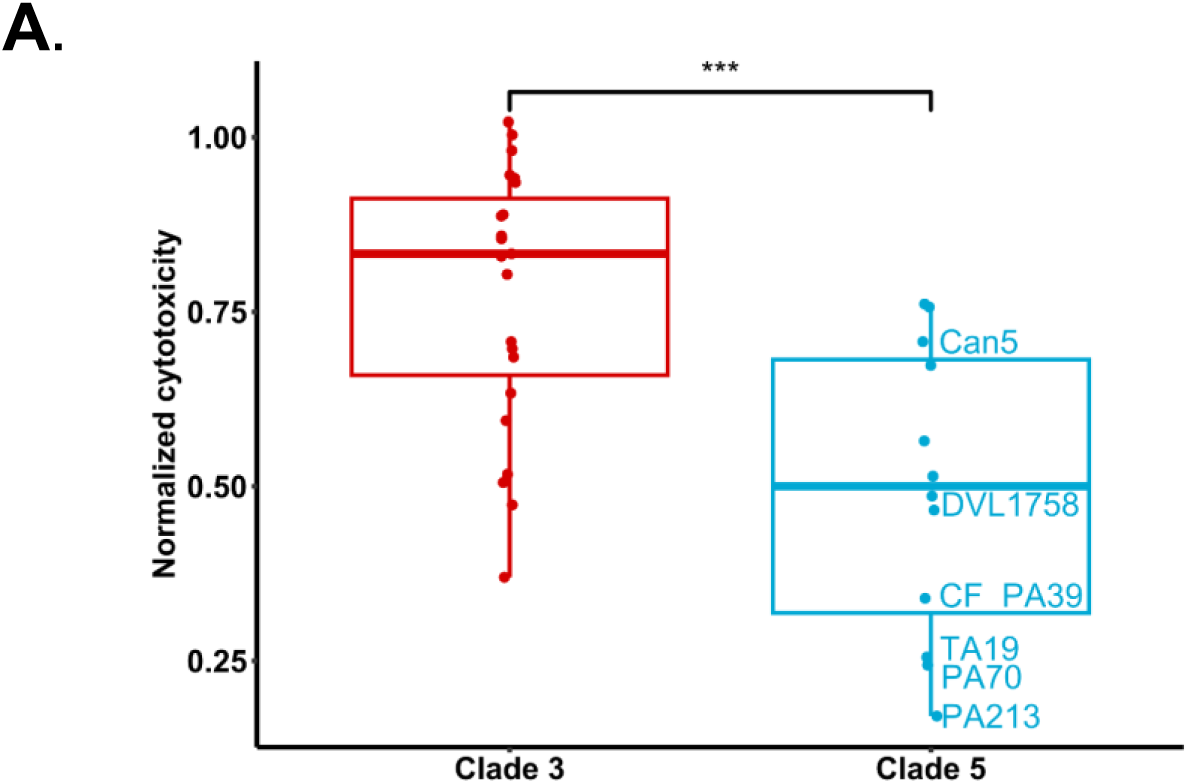
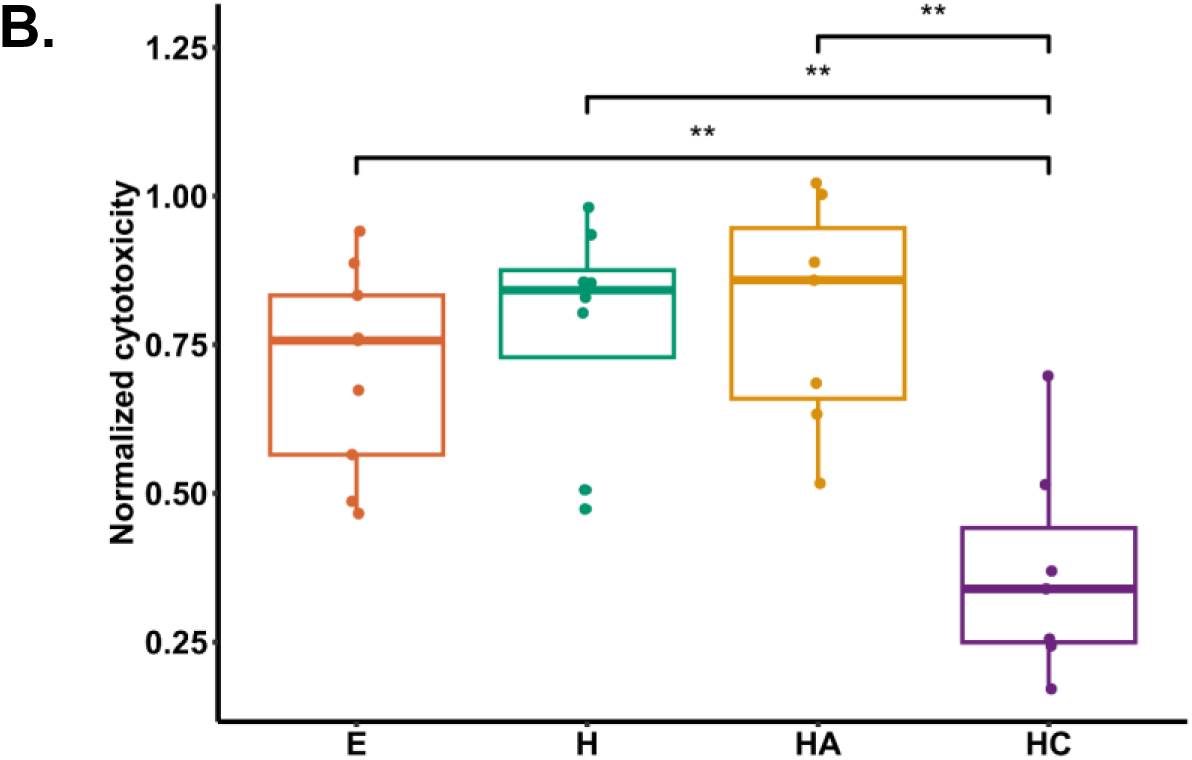
Strains from clade 5 and from chronic infections display lower cytotoxicity. Bacterial cytotoxicity on A549 epithelial cells measured by incorporation of propidium iodide. Each individual strain was assayed in at least two independent experiments and in triplicates (Figure S1), and area under the curve (AUC) were calculated from the obtained kinetics and normalized to a reference strain. **A**. Strains were grouped by clade and medians of AUC were calculated. The strains selected for further analysis (TA19, CF39, DVL1758, PA213, PA70 and Can5) are highlighted. **B.** Normalized cytotoxicity was analysed according to strains’ origin (E, environment; H, hospital; HA, acute infection; HC, chronic infection). The group of strains from chronic infections display statistically lower cytotoxicity compared to other three categories. **: p-value < 0.01 and ***: p-value < 0.001.

### Several strains from the clade 5 are avirulent in *Galleria* model of infection

We previously established the *Galleria mellonella* larvae model to determine the levels of virulence of ExlA-positive, T3SS-negative strains (Santausa et al., 2020). The kinetics of killing of *Galleria* due to injection of 5-10 colony forming units of *P. aeruginosa* correlated to the synthesis of ExlA (Trouillon et al., 2020).

Here we applied this model of infection to examine the virulence potential of all 35 strains of our collection (Figure 2B). Each strain was tested independently at least twice using twenty larvae per experiment (for details, see Material and Methods). As expected, the most cytotoxic strains *ex vivo* were also pathogenic to *Galleria,* killing all 20 larvae in >24h (JT87, IHMA87). Comparison of calculated AUC showed that most of the strains displayed intermediate pathogenicity. Some strains from both clades exhibited low or no virulence in this insect model with less than 10% of dead larvae after 24h: ATCC9027, AZPAE14941, WH-SGI-V-07618, WH-SGI-V-07370, WH-SGI-V-07234, WH-SGI-V-07064, WH-SGI-V-07050, (30 % of clade 3 strains) and CF_PA39, PA70, PA213, TA19, AL198 (42 % of clade 5 strains). In contrast, only strains from clade 3 killed more than 75% of the larvae after 24h: AZPAE14901, IHMA_879472, CPHL11451, JT87, EML528, WH-SGI-V-07072 and CLJ1 (30% of strains belonging to clade 3).

Surprisingly, some strains, despite their cytotoxicity *ex vivo*, had low pathogenicity profiles in *Galleria* (WH-SGI-V07370, WH-SGI-V-07618, ATCC9027), an observation made previously for ATCC9027 (Garcia-Reyes et al., 2021). The reasons for this discrepancy may be multiple, including the lack of specific components required for bacterial defense against the larvae immune system.

Globally, a correlation between cytotoxicity to A549 cells and virulence in insect larvae was observed with lower AUC of PI incorporation being associated with a higher percentage of survival after 24h of infection (Figure 3C). The strains considered avirulent in *G. mellonella* encompass two populations with different levels of cytotoxicity on cultured cells: strains with low to moderate cytotoxicity (from both clades) but also strains from clade 3 with high cytotoxicity (WH-SGI-V07050, WH-SGI-V07287, WH-SGI-V07370, WH-SGI-V07618 and ATCC9027). The lack of virulence of these cytotoxic strains remains to be investigated.

**Figure 3.**
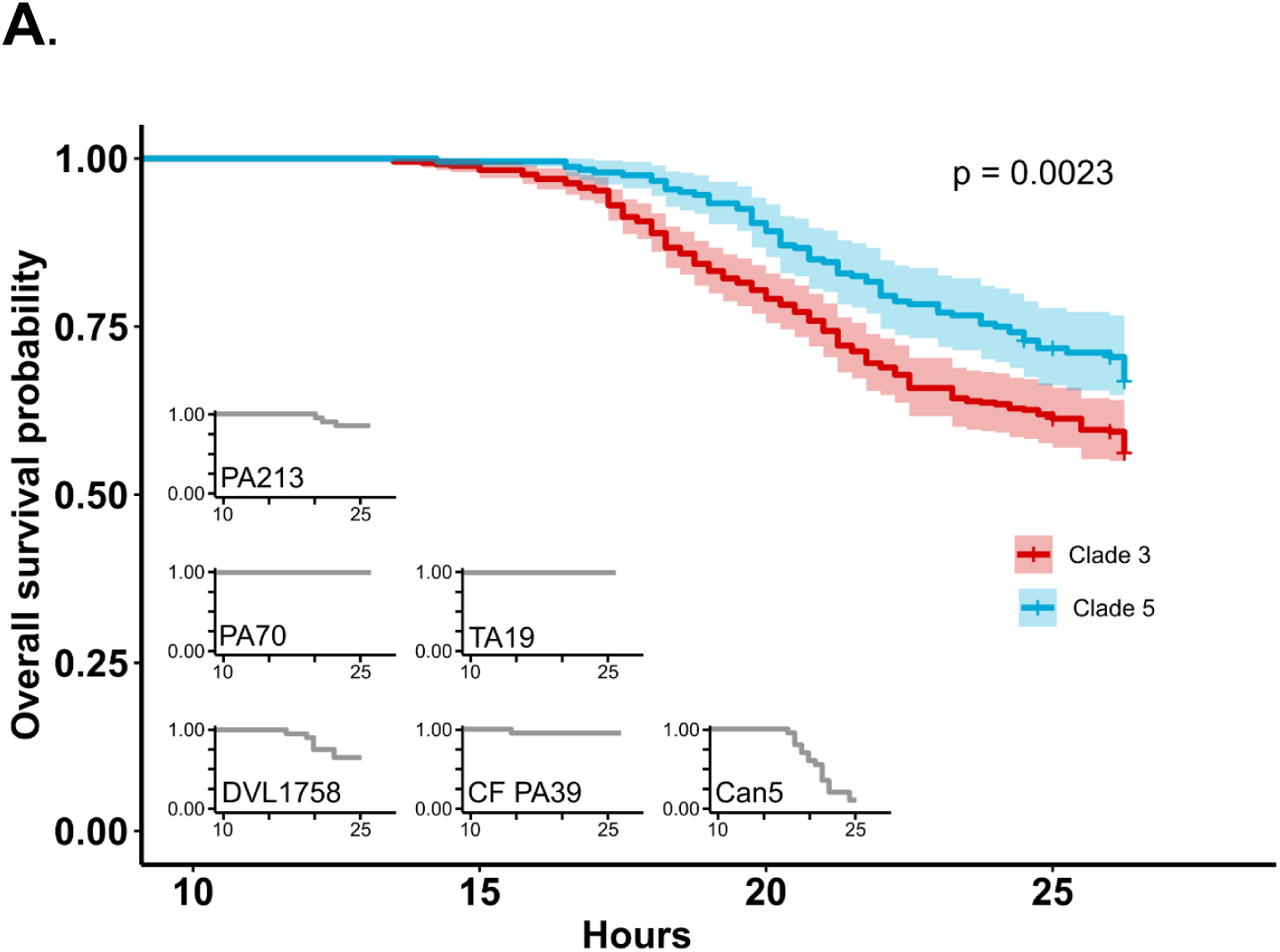
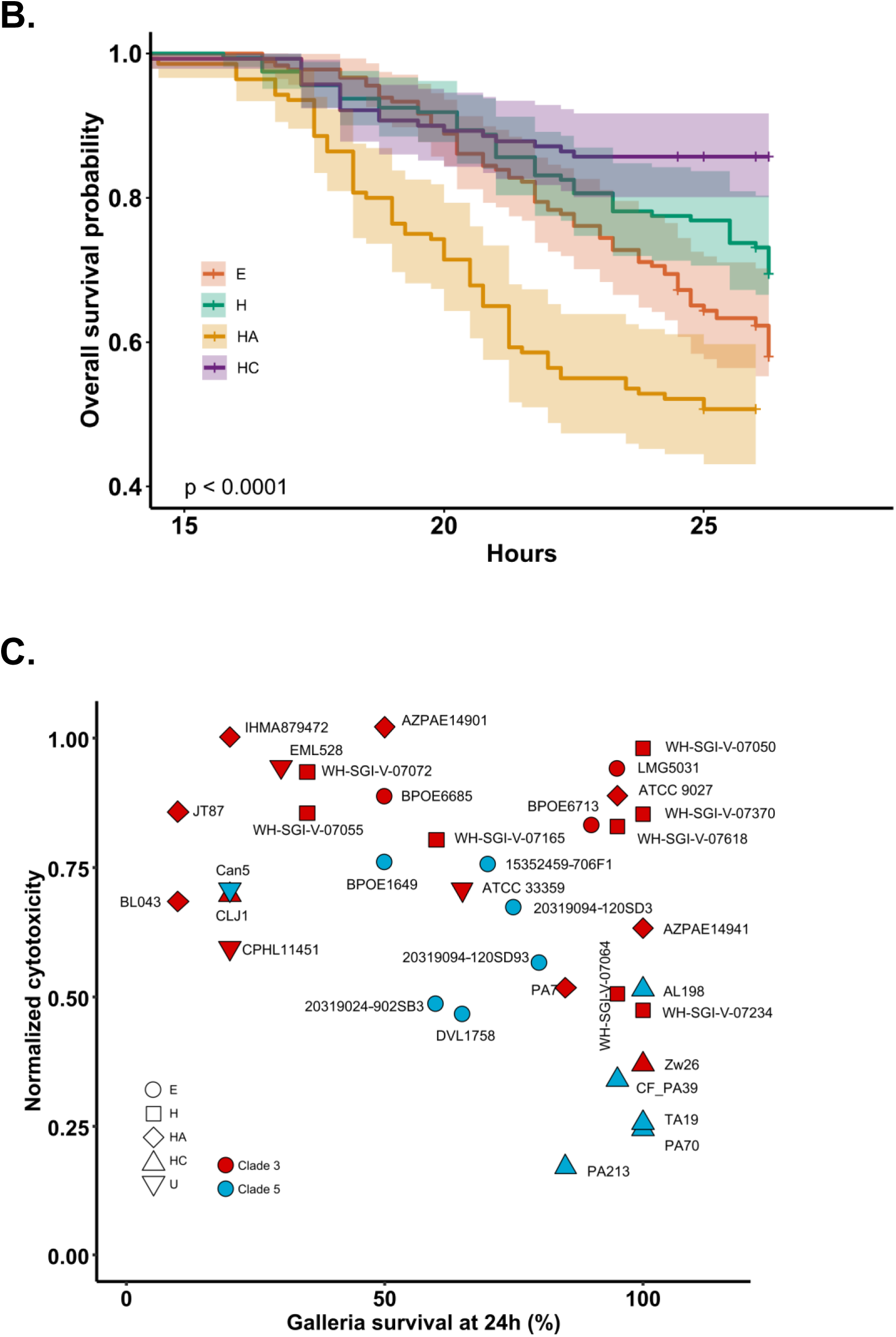
Virulence of strains assessed in *Galleria* model of infection reveals differences between clades and strain origins. **A**. The strains belonging to clade 5 show statistically (p=0.0023) lower virulence. The overall survival probability (one representative experiment) of strains selected for further analysis (TA19, CF39, DVL1758, PA213, PA70 and Can5) are displayed in insets. **B.** Strains grouped by origin (E, environment; H, hospital; HA, acute infection; HC, chronic infection) display gradual virulence profiles (HC>H>E>HC) that are all significantly different between each other (overall p-value <0.0001 and paired p-values < 0.05). **C.** Correlation between the cytotoxicity and virulence.

Finally, injection of the non-cytotoxic strains Zw26, PA70, TA19, and PA213 showed very limited larvae death, comparable to the levels of larvae injected with the buffer only, confirming the low virulence potential of those strains in the mice model of acute infection (Reboud et al., 2016).

The survival curves obtained with the different strains uncovered a significantly higher virulence of clade 3 compared to clade 5. Furthermore, considering their origins, the strains ranked into three groups with significant virulence differences: 1) strains from acute infections and the environment were the most virulent; 2) strains from undefined hospital origin showed an intermediate virulence while 3) the ones from chronic infection exhibited the lowest virulence.

### Strains of clade 5 are sensitive to common antibiotics

To determine the antibiotic resistance profile of our collection, each strain was tested against a panel of 16 antibiotics according to EUCAST 2018 standards. The diameter of growth inhibition and minimal inhibitory concentration were recorded. For each antibiotic and strain, the profile was expressed as RIS (Resistant, Increased exposure and Susceptible) and the score was calculated from R=2, I=1, S=0 (Figure 1, Table 1S) of all antibiotics tested. As expected, the recent hospital isolates had the highest resistant scores (up to 28 for WH-SGI-V-07165/VTK106158 and JT87), while environmental strains were sensible to most of the tested antibiotics. The strains were classified as sensitive to all tested antibiotics (S), non-<u>m</u>ulti<u>d</u>rug resistant (non-MDR), MDR and e<u>x</u>tensively (X)-DR. Several environmental strains were sensitive to all tested antibiotics, whereas one strain of the collection was found XDR (WH-SGI-V-07165). Therefore, the *exlA*^+^ collection contains *P. aeruginosa* isolates that are sensible to a variety of commonly used antibiotics, with several avirulent strains being susceptible to all antibiotics tested.

### Production of secondary metabolites in selected strains of clade 5

Given the high variability observed in ExlA-related phenotypes and the significance of secondary metabolite production in *P. aeruginosa*, we examined the production of three major secondary metabolites PYO, RL, PVD and the enzyme ELA in a subset of strains from clade 5. The strains selected for this analysis were PA39, PA70, DVL1758, PA213, TA19 and CAN5, based on their delayed cytotoxicity on epithelial cells and low virulence in *Galleria* assays.

No significant differences were observed in ELA production for strains PA39, PA70, and DVL1758, when compared to the reference strain PAO1. However, strain PA213 produced three times less ELA than PAO1, and strains CAN5 and TA19 showed no detectable ELA production (Figure 4A). For PVD, strains PA39 and DVL1758 produced similar levels to PAO1, indicating functional iron acquisition pathways. In contrast, strains PA213 and CAN5 produced three times less PVD, while strains PA70 and TA19 did not produce PVD at all (Figure 4B). Strain DVL1758 was notable for producing more PYO than the PAO1 reference, suggesting enhanced activity of the PYO biosynthesis pathway, potentially due to upregulation or mutations that favor PYO accumulation. No PYO was detected in the other strains (Figure 4C).

**Figure 4.**
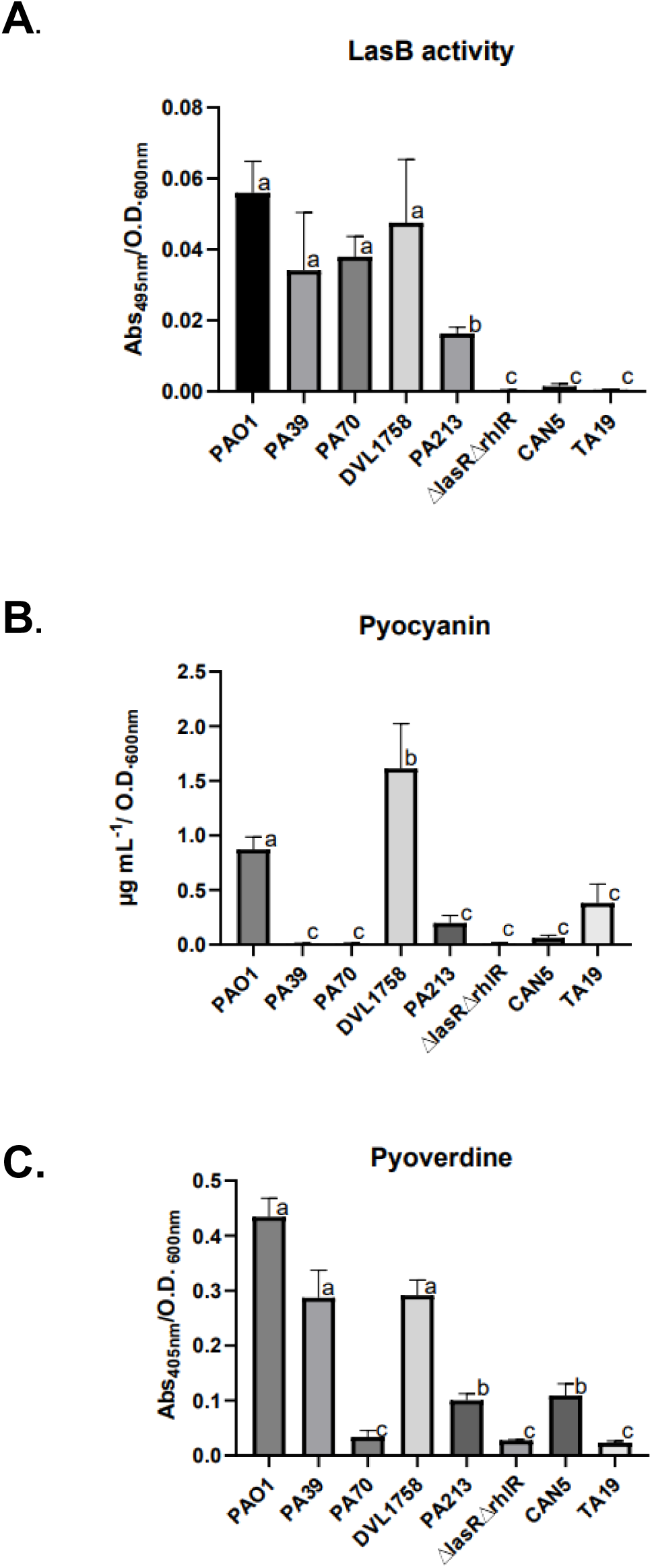
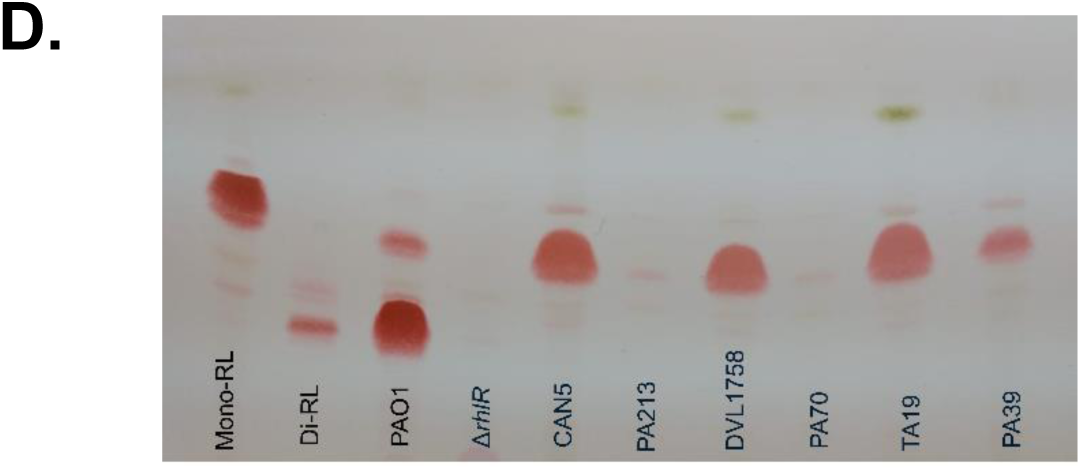
Production of LasB elastase (**A**) and secondary metabolites pyoverdine (**B**) and pyocyanin (**C**) by a subset of strains. The data of the three replicates were analyzed by Tukey test (a = 0.05), different letters indicate significant differences among the sample means; equal letters indicate no significant difference. (**D)** TLC detection of mono-RL and di-RL production in PPGAS media at 24 hrs of growth.

Finally, for the RL production, strains CAN5, DVL1758, and TA19 produced higher amounts of mono-RL than PAO1. Conversely, strains PA213, PA70, and PA39 showed low or undetectable levels of mono-RL (Figure 4D).

Previous studies have established the role of QS in the synthesis of secondary metabolites PVD, PYO and RL, as well as ELA, in *P. aeruginosa* (Gambello & Iglewski, 1991; Hall et al., 2016; Pearson et al., 1997; Dell’Anno et al., 2022). The different quantities of secondary products in strains tested here suggested the variability in QS functioning or regulation.

Two strains, CAN5 and TA19, were ELA- and PYO-negative but produced mono-RL on PPGAS media. The TA19 strain is a *lasR* mutant since it has a stop codon in the *lasR* gene (Figure S2), which explains its phenotype. However, the CAN5 strain has no defect in the Las, Rhl, or Pqs system, indicating that the observed phenotype depends on other mechanisms. We speculate that this strain has a defect in the Las system similarly to the *P. aeruginosa* ATCC9027 strain, which does not produce ELA and PYO in LB media but still produces mono-RL in PPGAS, because the Rhl system may operate independently of LasR in PPGAS (Grosso-Becerra et al., 2016)

PA70 and PA213 were negative for PYO and mono-RL but positive for ELA production. It is possible that in these strains, the Las system is active, with LasR acting as a negative regulator of pyocyanin, which allows for ELA production while suppressing PYO. Indeed, Soto-Aceves et al. reported the negative effect of LasR on PYO production in the PAO1 strain (Soto-Aceves et al., 2022).

The strain PA39, which was negative for PYO but positive for ELA and mono-RL, may have a defect that prevents the PqsE synthesis, similarly to the mutation in *pqsE* in the strain PAO1 (García-Reyes, Cocotl-Yañez, et al., 2021) The distinct regulation in PA39 suggests that the pathways controlling PYO and PVD production are separate, as this strain maintains normal PVD levels despite its inability to produce PYO. Interestingly, DVL1758 produced higher levels of PYO and mono-RL than the reference strain PAO1, along with ELA and PVD at comparable levels. Out of 25 strains of our collection, five carry deficient *lasB,* which - although less frequent then reported by Trottier et al., 2024 - confirms the occurrence of *lasB* mutants across strains of all clades and from different environments. We confirm also that the *lasB* mutations do not directly correlate with strains’ virulence and the level of production of secondary metabolites.

## Conclusions

In this work, we performed genetic and phenotypic analysis of a cohort of *P. aeruginosa* strains lacking the most recognized *P. aeruginosa* virulence factor, T3SS. We confirmed that 35 strains of our collection possess the *exlBA* operon, with potential of expressing the cytolytic toxin ExlA. However, despite the *exlBA* presence in the genomes, several strains were found non-cytotoxic on epithelial cells and avirulent in a *Galleria* model of infection, suggesting that the operon is not expressed under those conditions. Genome analysis highlighted additional specific signatures differentiating the clades 3 and 5. Finally, the scan of secondary metabolites and the quantification of the elastase enzyme in avirulent strains, showed highly variable levels of their production, suggesting the defects in the quorum regulatory system. Several strains (DVL1758, CAN5, TA19 and PA39) show potential for the industrial production of mono-RL. Strain PA70 could be valuable for producing ELA without PVD or PYO contamination, whereas PA39 could be used for PVD or ELA production without PYO interference.

## Acknowledgments

Special thanks go to Prof. Steve Lory, HMS, for his constant support and access to the sequencing facility, and to Peter Panchev for the *Galleria* experiments. *P. aeruginosa* IHMA87 was obtained from the International Health Management Association, USA. We would like to thank the *Pseudomonas* community to have provided other T3SS-negative *exlA*-positive strains. SG-R thanks Secretaría de Educación, Ciencia, Tecnología e Innovación de la Ciudad de México (SECTEI), for the scholarship awarded in the program for postdoctoral stays abroad.

The work was supported by the Laboratory of Excellence GRAL, financed within the University Grenoble Alpes graduate school (Ecoles Universitaires de Recherche) CBH-EUR-GS (ANR-17-EURE-0003) and the Fondation pour la Recherche Médicale (Team FRM 2017, DEQ20170336705) to I.A. Work in the CB laboratory is financed by the Agence National de la Recherche grant n°ANR-10-LABX-62-IBEID and the Fondation pour la Recherche Médicale (FRM) grant N° EQU201903007847

## Author contributions

IA designed the study and wrote the manuscript. SGR, MRG, YC and VCF performed experiments and analyzed the data, CR and LGV performed genomic analysis, EF and LG analyzed the data and prepared the Figures, CB, GSC and IA provided tools, supervised the work and validated the data. All authors edited the manuscript.

## Supplemental Figures and Tables

**Table S1.**
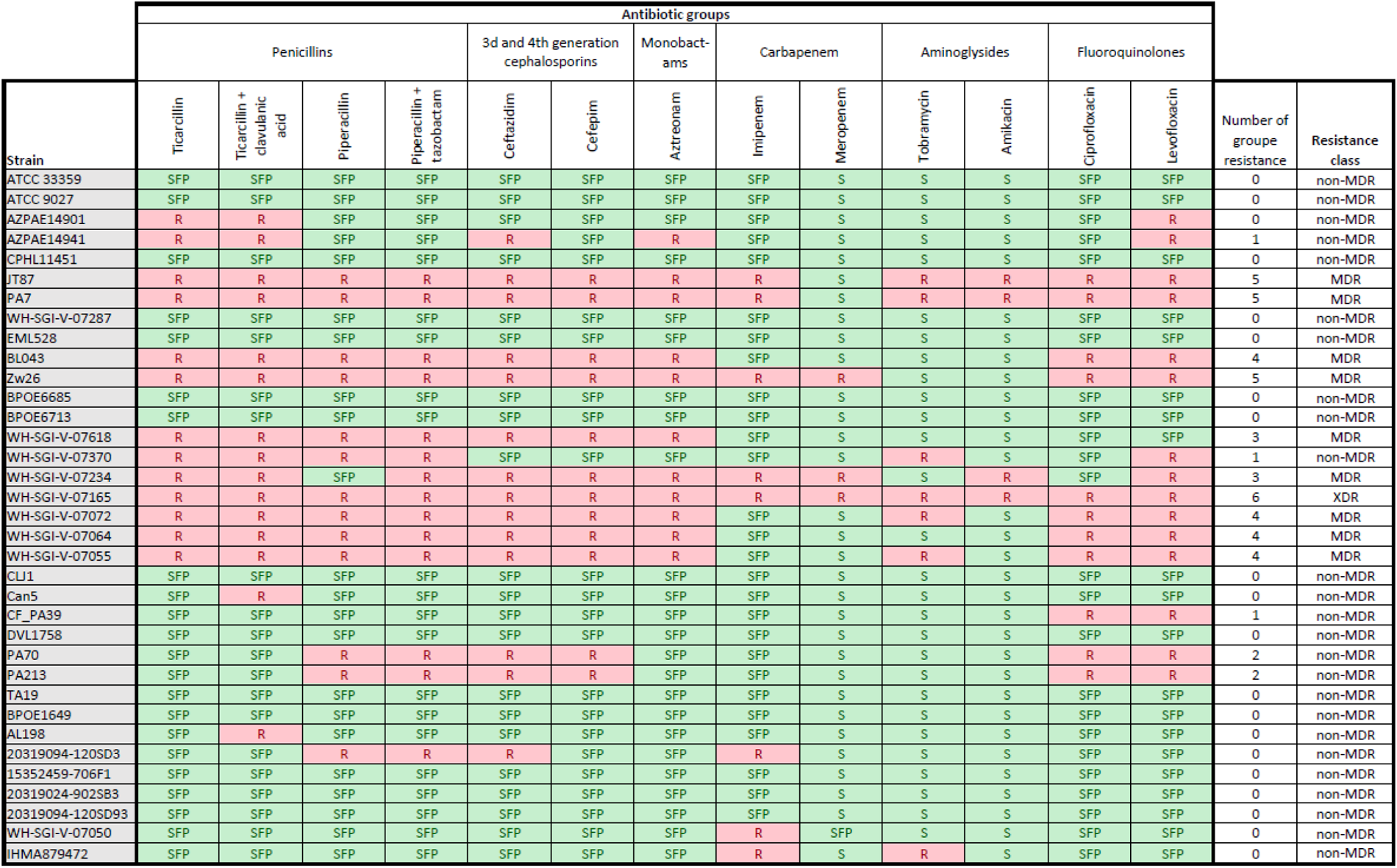
Antibiotic susceptibility tests on all strains of our collection. Resistance class is reported on Figure 1.

**Figure S1.**
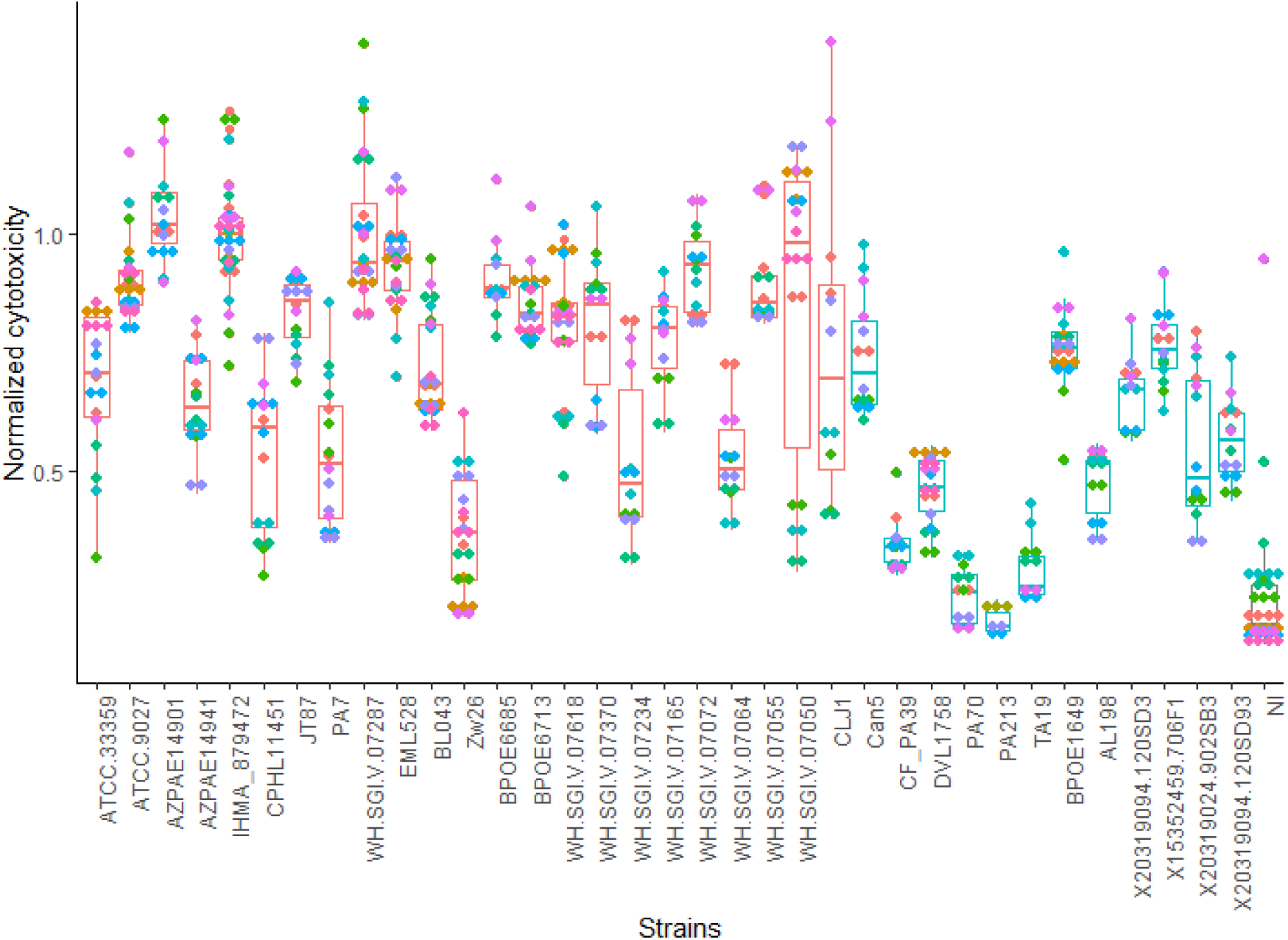
Cytotoxicity data. Each dot represents individual AUC. The triplicates from the same experiments are indicated with the same colors.

**Figure S2.**
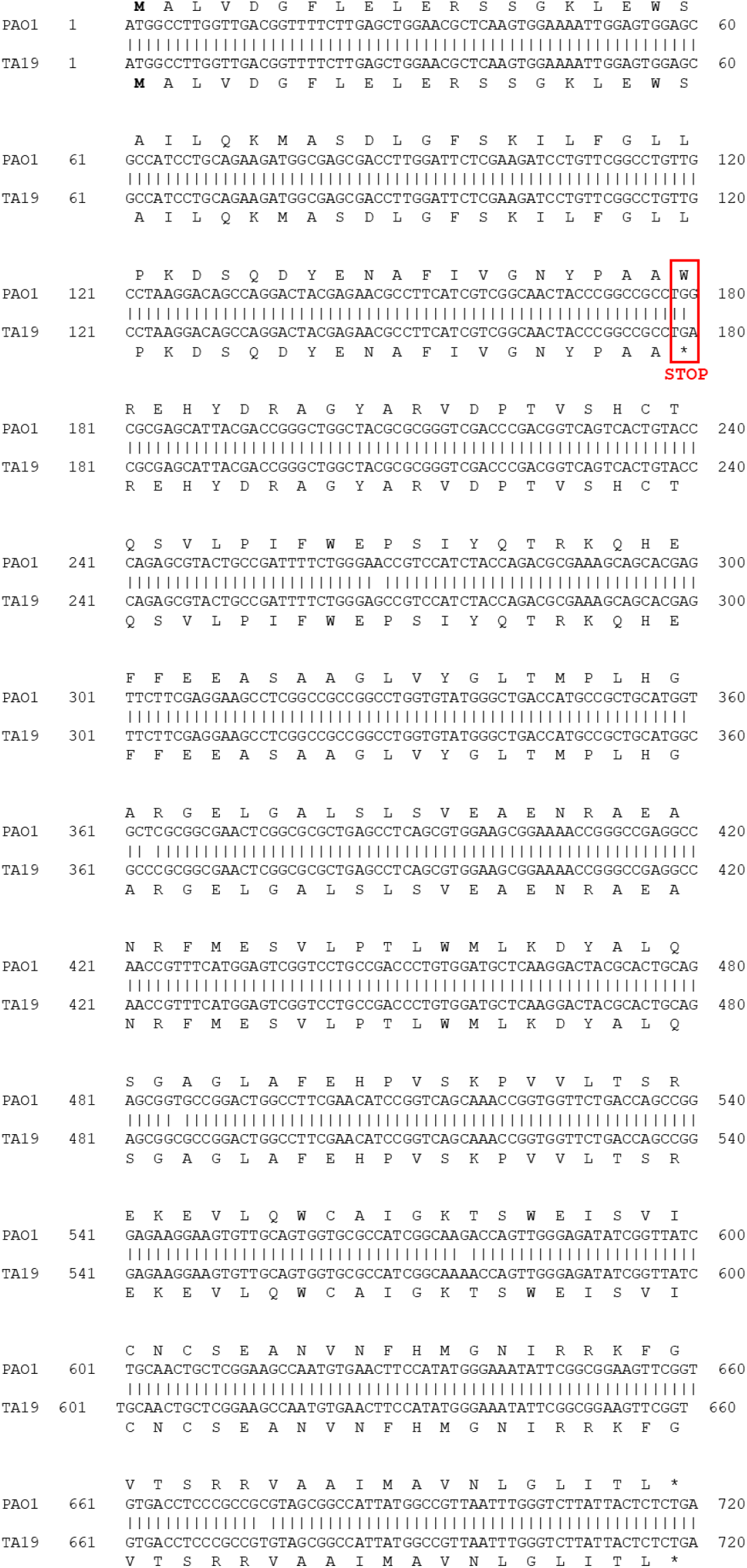
Global alignment of *lasR* gene of PAO1 and TA19 strains. The ATG is highlighted in bold and the site where the change from G to A occurs at position +180 in *lasR* TA19 that generates the stop codon is indicated in red. The protein sequence alignment was performed using the EMBOSS Needle tool available on the European Bioinformatics Institute (EBI) platform (https://www.ebi.ac.uk/jdispatcher/psa/emboss_needle).

